# Microscopic Control of Cortical Flows in Polarized *C. elegans* Zygotes

**DOI:** 10.1101/2025.01.12.632635

**Authors:** Elizabeth D. White, Younan Li, Rachel Kadzik, Yuqing Qiu, Aaron R. Dinner, Edwin M. Munro

## Abstract

Cell polarization, migration, and cytokinesis rely on flows of the cell cortex, a network of actin filaments, cross-linkers, and motors beneath the plasma membrane of animal cells. While actin network architecture, assembly dynamics, and motor activity are known to be important for cortical flows, how their modulation tunes macroscopic flow dynamics remains poorly quantified *in vivo*. Here, we use quantitative microscopy to constrain agent-based simulations that account for filament assembly, crosslinking, and motor activity. We calibrate the model to reproduce steady-state flows in polarized *C. elegans* zygotes and then challenge it to predict the results of RNA interference (RNAi) experiments. Our model predicts, and experiments largely confirm, a biphasic dependence of flow speed on microscopic rates of actin filament assembly and crosslinking. This biphasic dependence reflects a competition between the tendencies of perturbations to disrupt both transmission of and resistance to cortical forces. Our results provide new insights into how variations in microscopic features shape the emergent dynamics of the cell cortex. By establishing a well-calibrated model of cortical flow in a highly tractable model cell, we also provide a foundation for future studies of microscopic origins and biological control of cortical contractility and flow *in vivo*.

## Introduction

The cell cortex is a network of actin filaments, crosslinkers, myosin II motors, and other actin-binding proteins associated with the inner face of the plasma membrane of animal cells^1^. Forces generated within the cortex by myosin II drive cortical deformations and intracellular flows that underlie cell polarization^2–5^, cell division^6–10^, amoeboid cell migration, and tissue morphogenesis ^11–19^. Cortical flows require the continuous production, transmission and dissipation of forces within cortical actin networks. Each of these processes depends on network architecture, which is shaped by the interplay of many microscopic processes, including nucleation, elongation, capping, severing, and depolymerization of single filaments, as well as crosslinking of multiple filaments into higher order networks^20–23^. While these basic processes have been studied intensively, how they are integrated to shape network structure and dynamics in specific biological contexts remains poorly understood, owing in part to a lack of knowledge about molecular densities and rates *in vivo*.

Carefully calibrated simulations can in principle address this issue. Previous work has established several well-parameterized models of actin filaments, myosin motors, and other actin-binding proteins^24–27^ that perform well in predicting the structure and dynamics of actin networks reconstituted from purified proteins in vitro^28–30^. These and similar models have also been used to study contractile processes *in vivo*, such as contractile ring formation^31–33^, chromosome segregation^34^, and cortical flows^35^. These studies provide insights into how collective dynamics could emerge from microscopic interactions (known or hypothesized) and how structures and dynamics depend on molecular densities and reaction rates. However, comparisons between simulations and *in vivo* observations have remained primarily qualitative, largely because the simulations have not been grounded by measurements of microscopic properties in the system of interest. Here, we explore whether agent-based models tightly constrained by microscopic measurements in a particular experimental system can predict quantitative trends in both normal and experimentally perturbed dynamics observed *in vivo*.

To this end, we combine agent-based simulations and experiments to investigate how the microscopic dynamics of filament assembly, crosslinking, and motor activity shape cortical flows in the polarized one-cell *C. elegans* embryo (also known as the zygote). Zygotic polarity is established and maintained through advection of polarity proteins by cortical flows. Polarity is established during interphase by pulsatile cortical flows^2^ and then maintained through mitosis by quasi-steady (non-pulsatile) cortical flows^36^. Here we focus on the latter because their quasi-steady nature facilitates the use of microscopy and quantitative image analysis to constrain key parameters and make direct comparisons between measured and simulated patterns of flow. We use quantitative microscopy to constrain the microscopic dynamics of actin assembly, crosslinking, and motor activity in an agent-based model so as to reproduce rates and patterns of mitotic cortical flow observed *in vivo*. Then we challenge the model to predict the effects of depleting three highly conserved factors that tune actin network connectivity in different ways: formin/CYK-1^8,37,38^, profilin/PFN-1^38,39^, and plastin/PLST-1^40–42^, which control filament nucleation, elongation, and bundling, respectively. To compare simulations and experiments quantitatively, we use paired measurements to estimate the strengths of the perturbations. Our simulations predict, and experiments largely confirm, a biphasic dependence of flow speeds on the strengths of each perturbation, with more subtle variations across perturbations. Weak to moderate reductions in connectivity increase flow speeds, while strong reductions lead to network tearing and a decrease in flow speeds. We identify a feature of the dynamics that is common to all three perturbations at the transition between the two behaviors, and we interpret it to indicate that the transition reflects a competition between the effects of reducing network resistance and reducing force transmission as the connectivity decreases. Our results show how cortical connectivity in *C. elegans* embryos can be tuned to maximize flow speeds while limiting the effects of network tearing.

## Results

*Known biology motivates the elements of a microscopic model for cortical flow.* Our initial goal was to develop a predictive model of cortical flows in polarized *C. elegans* zygotes during mitosis, using the Cytosim framework for agent-based modeling of cytoskeletal networks (https://gitlab.com/f-nedelec/cytosim)^27,43,44^. During mitosis, the posterior cortex, where flows occur, is composed primarily of long, unbranched actin filaments that are nucleated by formin/CYK-1^8,37,38^ and elongated by profilin^38,39^. These filaments are organized into small bundles that are decorated by the crosslinking protein plastin/PLST-1, which is abundant in early embryos and is required for normal cortical network organization and force transmission during polarization and cytokinesis^40–42^. Contractile forces are produced by myosin II, which is organized into bipolar minifilaments with a few dozen motors each^45–47^. During mitosis, a spatial gradient of myosin II activity, shaped by its upstream activator CDC-42^36,48^ drives quasi steady-state cortical flow from the posterior pole towards the anterior.

These molecular considerations informed our choice of elements in the agent-based model, as we detail further in later sections. These included semi-flexible actin filaments that undergo nucleation, elongation, and disassembly; crosslinkers that bind reversibly to actin filaments and promote filament bundling; and two-headed motors that bind and translocate filaments with force-dependent kinetics. We deployed these elements within a two-dimensional simulation region of 16 µm by 20 µm, representing a medial region of the cortex in which a spatial gradient of myosin II drives anterior-directed cortical flows, bounded on either side by points of near zero flow (further details below). This choice of domain allowed us to impose simple boundary conditions and standardize comparisons between simulations and experiments.

As we describe further below, within this domain, we assumed spatially uniform actin assembly, disassembly, and crosslinking kinetics and introduced a gradient of motors that drove flow. We calibrated our simulations using previous measurements and new microscopy and quantitative image analysis to tune simulation parameters to reproduce successively measurements of actin filament assembly rates and lifetimes, bundle sizes, motor distributions, and flow speeds in wild type embryos. Finally, we validated our model through its ability to predict the effects of depleting plastin/PLST-1, formin/CYK-1, and profilin/PFN-1.

### Actin filaments undergo distinct phases of assembly and disassembly

We first focused on developing a quantitative microscopic description of actin filament assembly and disassembly dynamics. By tracking individual formin dimers in embryos expressing formin/CYK-1 endogenously tagged with GFP (CYK-1::GFP), Li and Munro previously showed that linear actin filaments assemble in the *C. elegans* cortex at ∼1.5 μm/s^8^; using single-molecule imaging of labeled actin subunits, Munro and co-workers previously showed that polymerized actin disassembles rapidly at the cortex, with a mean lifetime of ∼8 s^8,49,50^. However, these single-molecule observations did not reveal the pattern of disassembly of individual filaments, that is, whether filaments disassemble from their pointed ends to produce treadmilling dynamics or stochastically through severing and disassembly along the entire filament. Because these different modes of disassembly could produce different emergent dynamics of contractility and flow, we sought to distinguish them here.

To this end, we used embryos expressing a GFP-tagged form of plastin/PLST-1 (PLST-1::GFP) to visualize actin filament turnover dynamics *in vivo* (Figure 1A; in this and all other figures, the anterior pole of the zygote is oriented to the left). In these embryos, PLST-1::GFP decorates actin bundles, and dynamic assembly and disassembly of individual filaments within bundles gives rise to changes in patterns of fluorescence intensity^40^. Using fast near-TIRF microscopy, we observed bright streaks of PLST-1::GFP that elongate rapidly from one end, and then with a delay, shorten rapidly at the other end (Figure 1B; Supplementary Movie S1). Using kymography, we determined that these streaks elongate at approximately the same rate as individual growing filaments, as measured with CYK-1::GFP^8^. Because plastin presumably binds with higher avidity to bundled filaments than to isolated filaments^51^, we assume that these streaks reflect bundles formed *de novo* by two or more filaments growing simultaneously in parallel (see Supplementary Video 1). Kymography also revealed that plastin streaks elongate and shorten with identical speeds, with a characteristic delay of ∼9 s from the appearance of the plastin signal at a given position and its subsequent disappearance (Figure 1C). Because this delay is similar to the mean lifetime of polymerized actin measured by single-molecule imaging^8,49,50^, we infer that the disappearance of PLST-1::GFP signal reflects the disassembly of actin filaments from their pointed ends. Although a clear relationship between assembly and disassembly can only be inferred from cases where two or more filaments elongate together, here, we extrapolate from these observations to assume that all formin-assembled filaments undergo the same mode of dynamic treadmilling, in which actin subunits disassemble from pointed ends with a fixed delay of ∼8 s from their incorporation at the barbed end.

**Figure 1:**
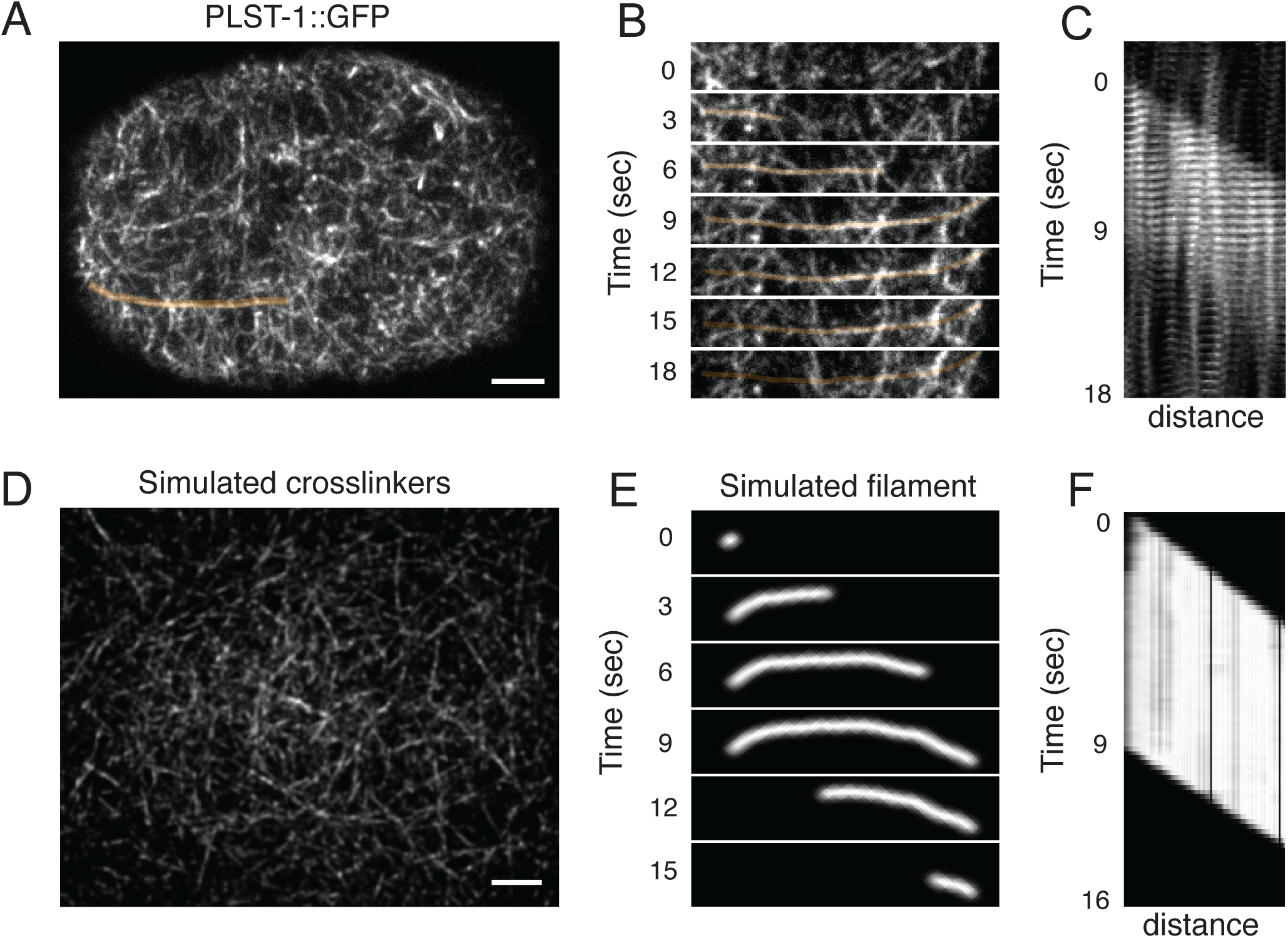
Actin filaments undergo dynamic treadmilling. (A) A polarized C. elegans zygote in late mitosis expressing plastin/PLST-1 tagged endogenously with GFP (PLST-1::GFP) Anterior is to the left in this and all subsequent micrographs. Rapidly elongating streaks of plastin (highlighted in orange) are consistent with the simultaneous assembly of two or more actin filaments by multiple formin dimers to form a small bundle. (B) Time course of the plastin streak (highlighted in A) illustrating progressive growth (right side) and progressive disassembly (left side) with a delay of ∼8 s. (C) Kymographs of five plastin streaks observed in one embryo, aligned with respect to their first appearance and averaged. (D) Pseudoimage of crosslinkers decorating the actin network obtained by convolving simulated data with a two-dimensional point spread function. (E) Pseudoimage of a simulated actin filament over its lifetime. (F) Kymograph of a simulated actin filament over its lifetime. Scale bars in (A) and (D): 5 µm.

To model these dynamics, we created a custom fiber class for Cytosim, representing actin filaments with three key behaviors: filaments nucleate at a fixed rate, they elongate from their barbed ends at a fixed rate for a variable duration before terminating growth, and they disassemble from their pointed ends (at the same rate as they assemble) with a fixed delay after nucleation (see Supplementary Movie S2). Based on our previous particle tracking analysis of formin/CYK-1 in the *C. elegans* zygote^8^, we set the actin filament elongation rate to 1.5 µm/s. In these experiments, the lengths of formin/CYK-1 trajectories often exceeded 10 µm, but more precise measurements of run length were limited by particle tracking errors and photobleaching. Thus, here we fixed the average duration for elongation to 8 s corresponding to a mean length of 12µm. We then tuned the nucleation rate to reproduce the overall density of filaments in simulations (see below). To facilitate comparison between the simulations and experiments, we convolved images from simulations with an approximated two-dimensional point spread function representative of our microscope (Figures 1D and 1E, see Methods). We were able to obtain good correspondence between experimental and simulated kymographs (Figures 1C and 1F).

### Actin filaments preferentially assemble along existing filaments and form bundles of various sizes

Having reproduced the dynamics of isolated actin filaments, we next sought to quantify the dynamics and extent of actin filament bundling to incorporate these dynamics into the simulations. Li and Munro previously observed that bundling occurs through a process termed filament-guided filament assembly in which growing filaments encounter and grow along existing filaments and bundles^8^. This process presumably reflects rapid binding of crosslinkers. The observation that plastin streaks grow at the same rate as individual filaments (Figure 1) suggests that plastin is sufficiently abundant that its binding rate is limited by the availability of proximally growing actin filaments. Thus, we assumed that filament-guided filament assembly is directed by the zippering of growing filaments onto existing ones, mediated by rapid plastin binding. Using a fixed unbinding rate of 0.05 s^−1^ ^40^, we found that a density of ∼70 μm^−2^ crosslinkers and a binding rate of 60 s^−1^ were sufficient to reproduce the experimentally observed filament-guided filament assembly in our simulations (Figure 2A). Pseudoimages of crosslinkers parameterized as described above closely resemble micrographs of *C. elegans* zygotes expressing PLST-1::GFP (Figure 1D).

**Figure 2:**
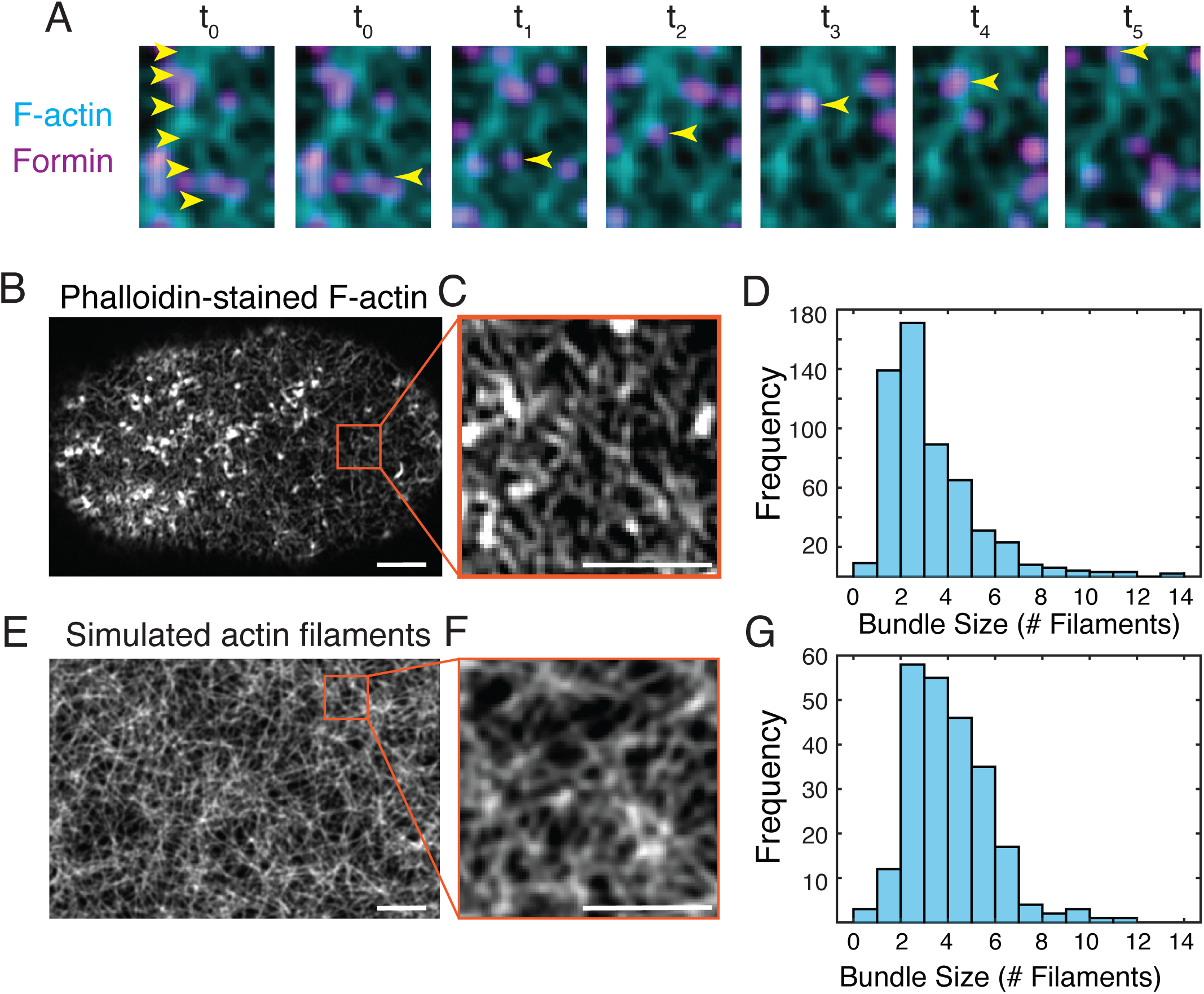
Actin filaments preferentially assemble along existing filaments and form bundles of varying sizes. (A) Pseudoimage showing elongating filaments (barbed ends marked by formin dimers (magenta) within a network of existing filaments/bundles (cyan). Yellow arrowheads in leftmost panel mark an existing filament bundle. Yellow arrowheads in subsequent panels (t0 – t5) mark the path of a formin at the tip of one filament elongating along an existing bundle. (B) A C. elegans zygote in mitosis fixed and stained with phalloidin to label F-actin. (C) Inset of (B), highlighted in red box, showing actin filament bundles with different local brightnesses, indicating variation in bundle size. (D) Histogram of bundle sizes in experimental images. (E) Pseudoimage of simulated actin filaments in simulation. (F) Inset of (E) highlighted in red box, showing actin bundles of different widths and brightnesses. (G) Histogram of bundle sizes measured in pseudoimages. Scale bars in (B) and (E): 10 μm. Scale bars in (C) and (F): 5 μm.

Finally, having constrained the assembly dynamics of single filaments, and the dynamics of local filament bundling, we tuned the filament nucleation rate so that simulations reproduced the distribution of actin bundle sizes observed in the posterior cortex during mitosis. To measure the distribution of actin bundle sizes, we fixed and stained mitosis-staged *C. elegans* embryos with phalloidin and then imaged them using laser scanning confocal microscopy with an AiryScan detector to produce high-resolution images in which individual bundles could be distinguished (Figures 2 A, B, and C). Because phalloidin binds stoichiometrically to polymerized actin subunits, the local brightness and width of individual bundles should be proportional to the number of filaments they contain. Therefore, following subtraction of the background intensity, we took transects across many individual bundles and used the integrated intensity under individual peaks along the transect as a proxy for bundle size. Assuming the smallest intensity values represent single filaments (see Methods for further details and discussion), the peak of the distribution is 3 filaments, and the average bundle size is 4 filaments (Figure 2D).

Finally, fixing all other parameters for filament assembly and crosslinking as described above, we varied action nucleation rates, allowed simulated networks to reach steady state, created pseudoimages (see Methods for further details), and used the same transect method to determine the simulated bundle size distribution (Figure 2 E and F and Figure 9). We found that a nucleation rate of 121 s^-1^ reproduced the bundle size distribution observed *in vivo* (Figure 2G).

### Reproducing cortical flow observed during maintenance phase

As discussed above, cortical flow is driven by a gradient of myosin II, which is patterned by the small GTPase CDC-42^36,48^. To quantify the relationship between the myosin gradient and flows for comparison to simulations, we imaged embryos expressing myosin II endogenously tagged with mKate2 (NMY2::mKate2^52^) at 1 s intervals during mitosis (Figure 3A and Supplementary Movie S3). For each embryo, we selected a time window of 60-80 s in mid-late mitosis, during which myosin densities and flow patterns were approximately steady. Within this time window, we computed the time-averaged axial intensity profile of myosin II (Figure 3B). We found that myosin II is clearly enriched in an anterior domain spanning ∼40% of the embryo’s length. Beyond the posterior edge of this domain, myosin II decays smoothly, reaching background levels near the posterior pole. Within the same time window, we computed the time-averaged axial flow velocity field by particle image velocimetry (PIV) using the Quantitative Fluorescence Speckle Microscopy (QFSM) software package developed by the Danuser group^53^ (Figure 3C; by our convention, negative velocity corresponds to anterior-directed flows). Anterior-directed flows reached a maximum speed in the medial region where myosin II forms an intensity gradient. Flow speeds approached zero, sometimes reversing direction, near the posterior pole, and near the posterior edge of the anterior domain, where myosin II is enriched. In all embryos, we observed a clear inflection point in the velocity curve near the posterior edge of the anterior domain. Using this inflection point to align flow profiles from different embryos, we found that they had similar maximum flow speeds and shapes (Figure 3D; see Methods for details). These observations define a medial region for direct comparison to simulations (marked by the red box in Figure 3, A-C) in which a spatial gradient of myosin II drives anterior-directed cortical flows, bounded on either side by points of near zero flow.

**Figure 3:**
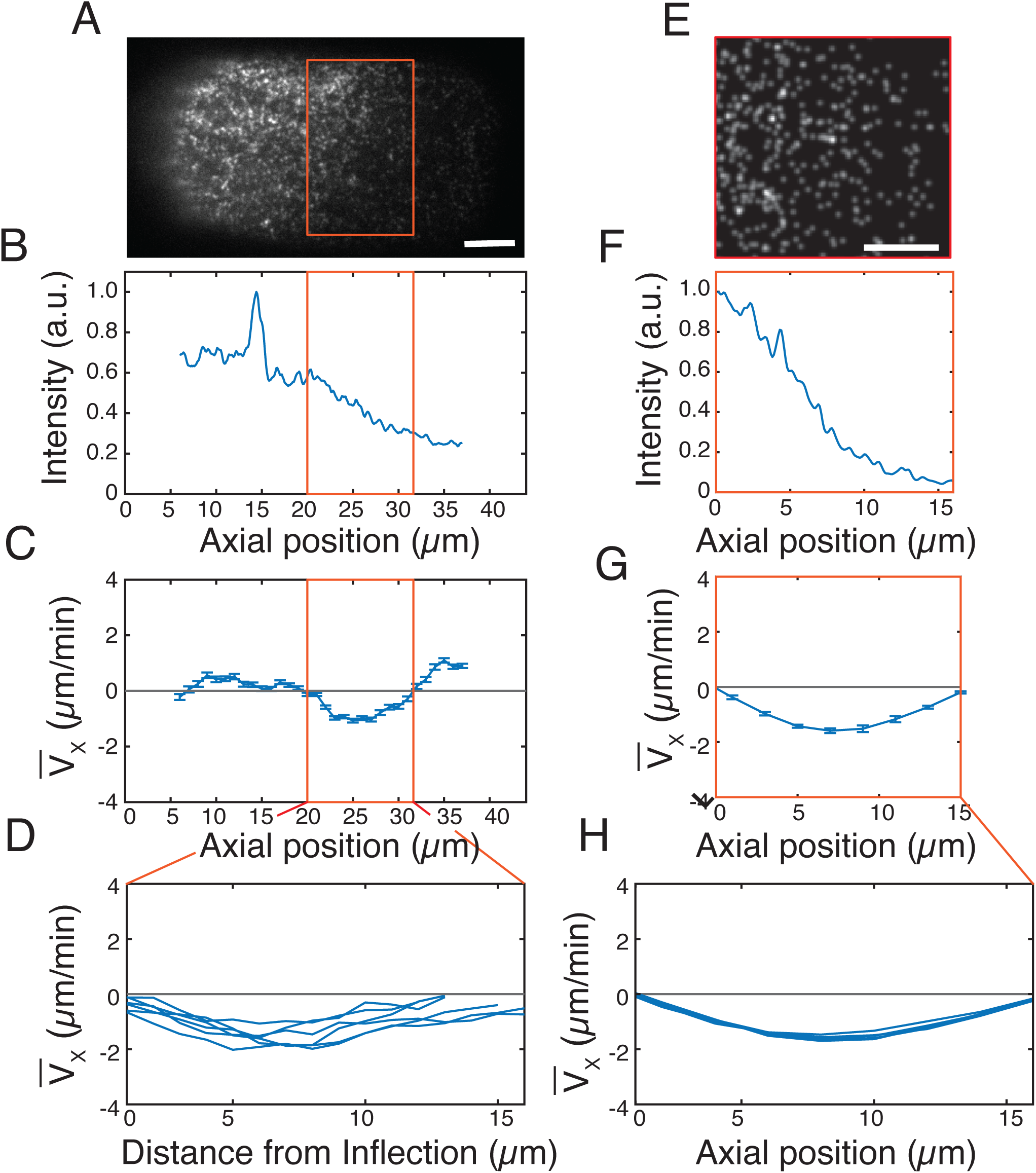
A myosin gradient drives cortical flow. (A) Surface view of a C. elegans zygote in mitosis expressing NMY2::mKate2. (B) Intensity profile of NMY2::mKate2 along the anterior-posterior axis, averaged over 60 frames. (C) Axial velocity (V_X_) vs position along the anterior-posterior axis, averaged over 60 frames. An inflection point in the velocity profile (left boundary of red box in (A-C)), present in all embryos, defines the anterior boundary of the posterior domain considered in simulations. (D) Axial velocity profiles for multiple embryos (n=6), aligned with respect to the inflection point. (E) Pseudoimage of steady-state myosin II distributions from simulations. (F) Axial intensity profile of myosin II from simulations, averaged over 60 s. (G) Axial velocity profile for a single simulation, averaged over 60 s. (H) Axial velocity profiles for 5 different simulations. Scale bars in (A) and (E): 5 μm.

Following previous work^27,32^, we modeled minifilaments of myosin II in Cytosim as two-headed bipolar motors. We endowed each motor head with a linear force-velocity relation as follows. Using our previously developed methods to track myosin II speckles on the posterior cortex of embryos during mitosis (see methods), we identified a subset of myosin speckles that appeared to move ballistically. Based on their average speeds, we set the unloaded speed of simulated motor heads to be 0.3 μm/s. We estimated the stall force to be the average number of myosin heads in a minifilament (10-20)^46,47^ times the stall force of a single myosin motor (∼3.4 pN)^54^ and thus set the stall force to 45 pN.

Having defined the properties of the motors, we tuned their rates of addition in different simulation regions (with fixed uniform removal rates; see Methods for details) to reproduce the steady-state myosin gradients and flow profiles observed in the medial zone (Figure 3 and Supplementary Movie S4). These results confirm that an agent-based model strongly constrained by microscopic observations of actomyosin organization and assembly dynamics can reproduce macroscopic dynamics of cortical flow observed *in vivo*.

### Simulations predict nonmonotonic changes in maximum flow speed in response to perturbations

We sought to test our model by comparing simulated and experimentally observed responses to perturbations without further parameter tuning. We focused on three experimental perturbations that we could map quantitatively to parameters of the model: depletion of formin/CYK-1, profilin/PFN-1, and plastin/PLST-1 by RNA interference (RNAi). These correspond to reduction of the filament nucleation rate, the filament elongation rate, and the crosslinker density, respectively. All three perturbations should decrease the connectivity of the network—the crosslinker density directly, and the actin nucleation and elongation rates through their effects on the mean length and density of actin filaments. The decrease in connectivity should reduce force generation and transmission, but it should also decrease viscous resistance; it can also lead to restructuring of the network and in turn more subtle effects. Depending on which effect is dominant, the flow speed could increase or decrease as the strength of a perturbation varies.

We quantified the effects of the perturbations by measuring the maximum time-averaged axial flow speed (hereon 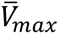). In all three cases, our simulations predicted a nonmonotonic trend in which moderate perturbations caused 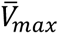 to increase, while sufficiently strong perturbations caused it to decrease (Figure 4, A, C, and E; yellow symbols indicate the parameter values used for unperturbed embryos). The transition between the increase and decrease (i.e., the peak in 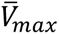) occurred at 30% of the unperturbed crosslinker density, 20% of the unperturbed actin nucleation rate, and 50% of the unperturbed actin elongation rate.

### Depletion experiments validate model predictions

As noted above, we used RNAi to deplete plastin/PLST-1, formin/CYK-1, and profilin/PFN-1 to evaluate these predictions experimentally. In each case, we varied the strength of the perturbation by varying the duration of exposure to RNAi, and we then made paired measurements in individual embryos of the strength of perturbation and 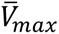 during mitosis. For depletion of plastin/PLST-1, we used a strain co-expressing endogenously-tagged myosin II (NMY-2::mKate2) and PLST-1::GFP, and we measured the strength of depletion by quantifying the density of cortical PLST-1::GFP during mitosis. For depletions of formin/CYK-1 and profilin/PFN-1, we used a strain co-expressing NMY-2::mKate2 and CYK-1::GFP. To quantify formin depletion, we quantified flows during mitosis as described above. Then in the same embryos, at the two-cell stage, we quantified the density of cortical formin dimers during mitosis, relative to wild type controls, and used the decrease in the density of formin speckles as a proxy for the decrease in the rate of actin filament nucleation. Similarly, we used the decrease in the speed of rapidly moving formin dimers, measured during mitosis in two-cell embryos, as a proxy for the decrease in the filament elongation rate in profilin-depleted embryos co-expressing NMY-2::mKate2 and CYK-1::GFP (see Methods).

Plots in Figure 4, B, D, and F show paired measurements of depletion strength and 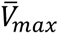 in mitosis for the three different perturbations, while kymographs in Figure 5 show representative axial patterns of cortical flow observed during mitosis in wild type embryos (Figure 5A) and in embryos with weak, moderate, and strong levels of depletion of profilin/PFN-1 (Figure 5B), formin/CYK-1 (Figure 5C), and plastin/PLST-1 (Figure 5D; see also Supplementary Movies S5-S7). Consistent with simulations, moderate depletions of profilin, formin, and plastin all caused increases in the rates of anterior directed flow (Figure 5, B-D, leftmost two columns, yellow arrows). Paired measurements of depletion strength and 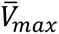 confirmed both the qualitative trend and the overall magnitude of the effect of moderate depletions on axial flow speeds (Figure 5, D-F).

**Figure 4:**
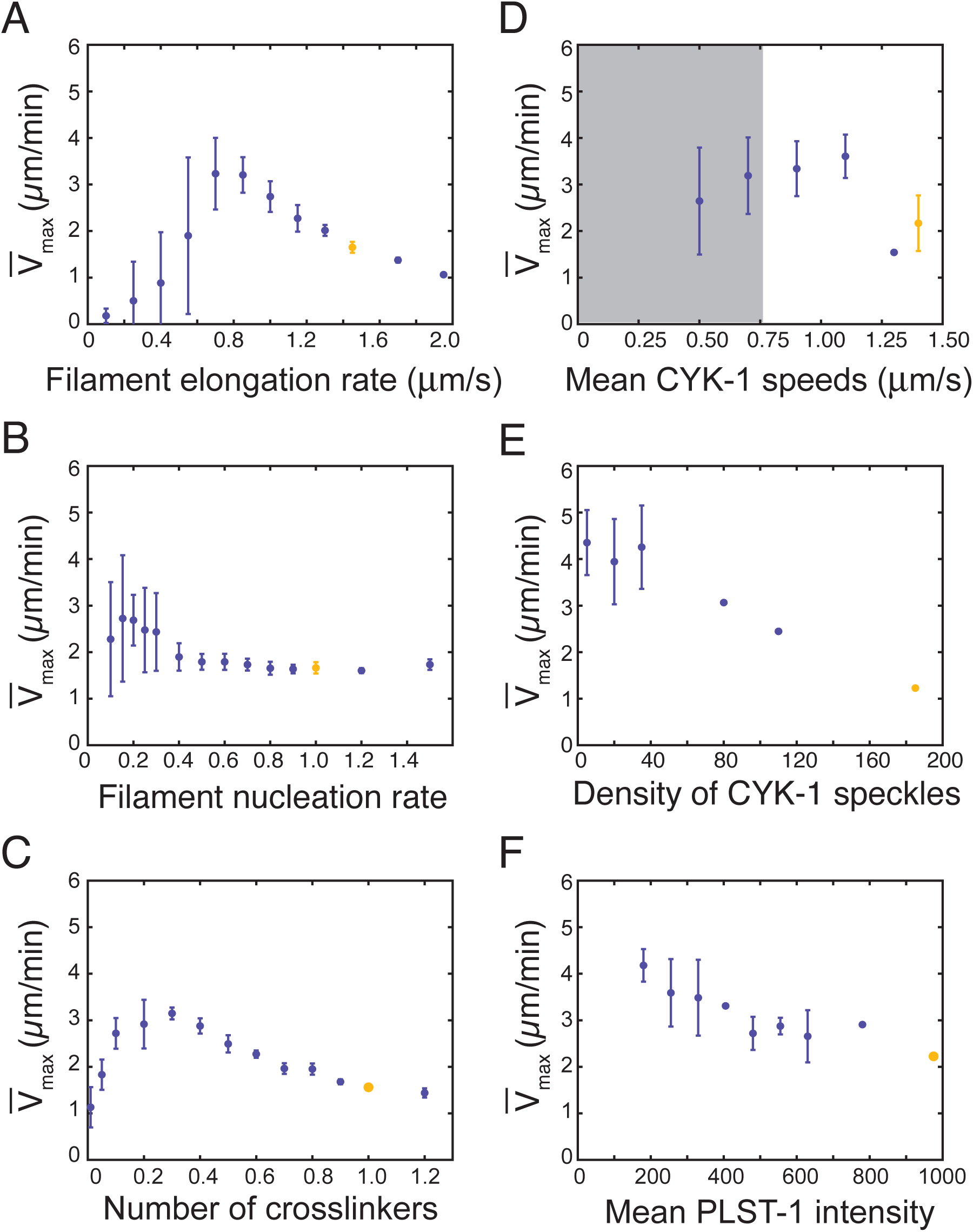
Simulations predict changes in maximum flow speed in response to perturbing plastin, formin, and profilin. (A,B) Maximum axial flow speed V_max_ as a function of crosslinker density relative to unperturbed conditions for simulations (A) and experiments (B). (C,D) V_max_ as a function of filament nucleation rate relative to unperturbed conditions for simulations (C) and experiments (D). (E,F) V_max_ as a function of filament elongation rate for simulations (E) and experiments (F). Yellow symbols correspond to unperturbed embryos. Gray box in (F) denotes where emeasurements of actin elongation rate are confounded by fast cortical movements caused by local tearing of the cortical network. Results for simulations are averaged over five simulations. In experiments, we computed V_max_ for individual embryos, we binned the data with respect to paired measurements of perturbation strength, and then computed the means and SEMs for each bin. (plst-1(RNAi): n = 27 embryos, pfn-1(RNAi): n = 22 embryos, cyk-1(RNAi): n = 33 embryos).

**Figure 5:**
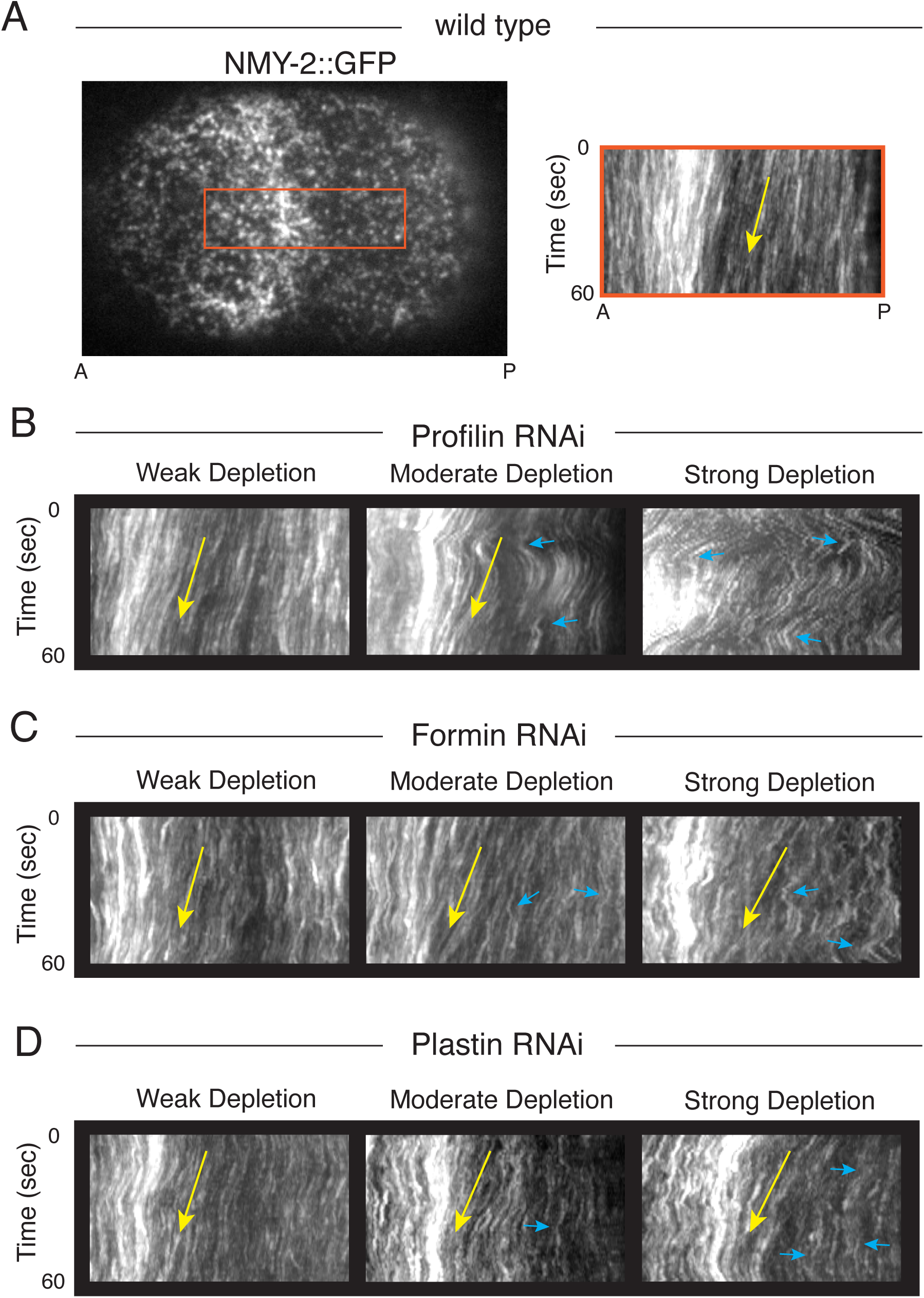
Depleting plastin, formin, and profilin alters patterns of macroscopic flow in distinct ways. (A) C. elegans embryo in mitosis expressing NMY2::GFP. Red box indicates the region used to create the kymograph shown to the right. (B-D) Kymographs of weak (left), moderate (middle), and strong (right) depletions for plastin/PLST-1 (B), formin/CYK-1 (C), and profilin/PFN-1 (D). Yellow arrows highlight flow towards the anterior pole. Blue triangles highlight local fluctuations in cortical flow speed and direction that correspond to local tearing events.

In the simulations, 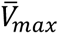 exhibited the greatest sensitivity to the filament elongation rate (i.e., the peak in Figure 4A is right of those in Figure 4, B and C). Consistent with this trend in the model’s predictions, moderate depletions of profilin/PFN-1 led to a peak in 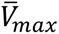 and strong depletions led to a decrease in 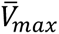. For these strong depletions, we observed numerous widespread cortical tearing events, consistent with loss of cortical connectivity; these appear in kymographs as abrupt changes in cortical flow direction and velocity (Figure 5B, rightmost panel, blue triangles; see also Supplementary Movie S5). We observed similar signs of cortical tearing with moderate depletion, but these were fewer and far less pronounced (Figure 5B, middle panel, blue triangles; see also Supplementary Movie S5).

In the simulations, the peak in 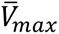 for the filament nucleation rate is left of and less pronounced than that for the filament elongation rate. Overall, the effect of depletion of formin/CYK-1 is consistent with this prediction, though no clear decrease is seen for strong perturbations. Similarly to the depletion of profilin/PFN-1, we observed a loss of coherence and local cortical tearing events, whose frequency and magnitudes increased with level of depletion (Figure 5C, blue triangles; see also Supplementary Movie S6); however, there were fewer such events for a given strength of depletion, consistent with the trends in maximum axial flow speed.

For the plastin/PLST-1 depletion, we did not observe a peak in maximum average flow speed, and we observed only a very mild loss of coherence, which appeared as reduced straightness of individual streaks within kymographs (Figure 5D blue triangles; see also Supplementary Movie S7). This may be due to incomplete depletion of PLST-1 by RNAi or the presence of other crosslinkers in the early embryo^42,55,56^. Similar considerations may account for the lack of a clear decrease in 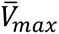 for strong depletion of formin/CYK-1^57^. Given these experimental uncertainties, the simulations and experiments are in remarkable quantitative agreement.

### Simulations connect cortical flow speeds and network dynamics quantitatively

Because there was no obvious network tearing in the simulations, we wanted to determine if a loss of coherence of the motion was responsible for the decrease in maximum axial flow speed for strong perturbations. The cortical tearing events that we observe experimentally were characterized by seemingly random motion in the direction perpendicular to the anterior-posterior axis; based on this, we computed the correlation of the velocity component in this direction as a function of distance for the simulations. The resulting correlation functions were well fit by an exponential form, and we thus characterized each condition by a single correlation length (Figure 6 and S4 and S5; see Methods for further details). Consistent with the greater sensitivity to reduction of the filament elongation rate than to the other perturbations, the former leads to shorter correlation lengths. Strikingly, however, the peaks in cortical flow speed for simulations shown in Figure 4 occurred in all cases when the correlation length was about 0.85 µm (we note that the density of actin scales linearly with the relative degree of depletion in the case of the filament nucleation and elongation rates). This suggests that, even though loss of coherence and tearing is not apparent in the simulations, the same physics underlies the biphasic behavior observed in simulations and experiments.

**Figure 6:**
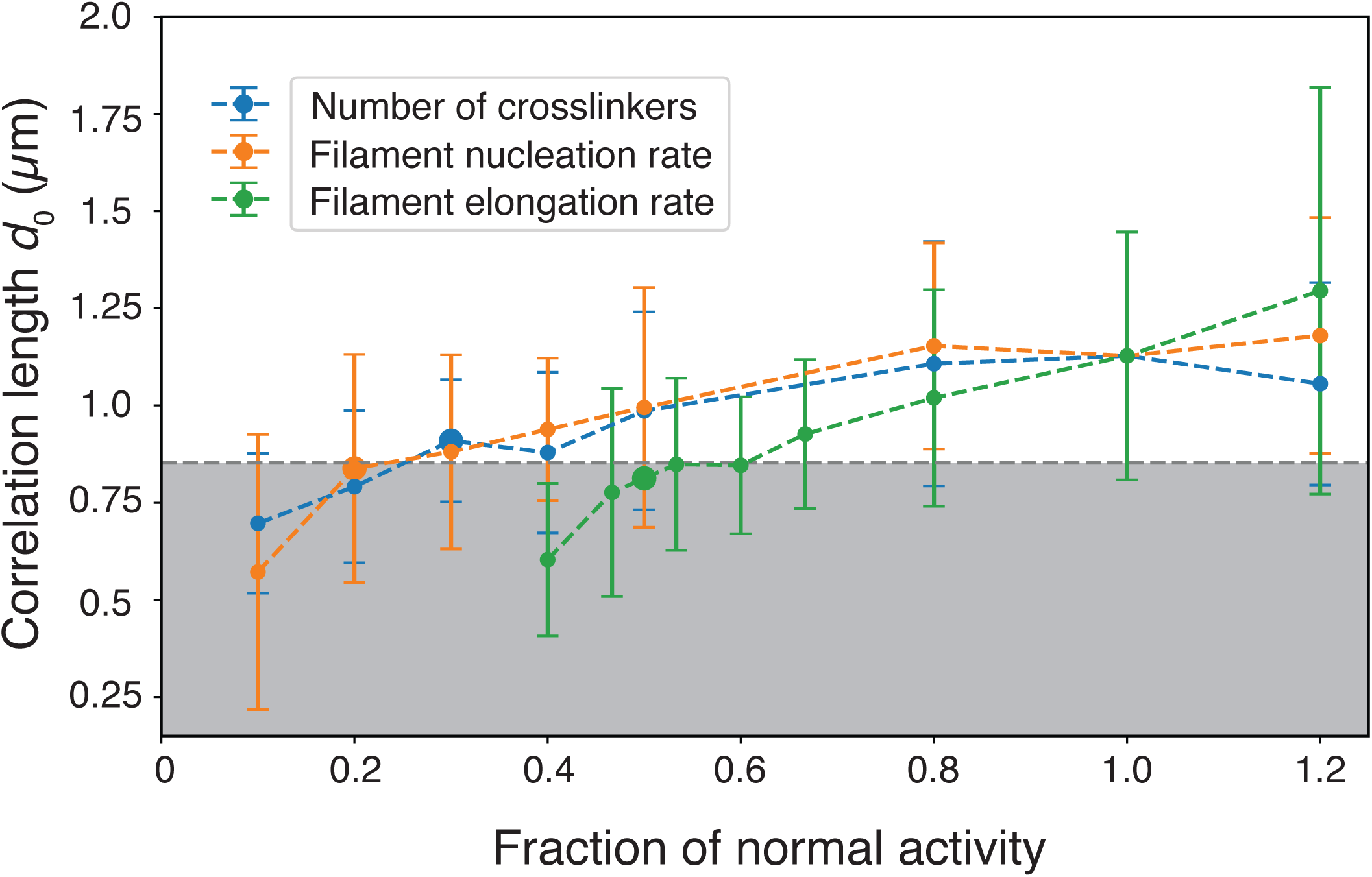
Correlation length of the velocity perpendicular to the anterior-posterior axis decreases as perturbation strength increases. Correlation lengths plotted as a function of fractional reduction in parameter values relative to their unperturbed values. Results shown are mean (symbols) and standard deviation (error bars) for 40 fits from 10 independent simulations for each condition (see Methods). Large circles mark the conditions at which the maximum axial flow speeds are achieved.

## Discussion

Here, we used a combination of simulations and experiments to explore how factors that regulate actin network architecture tune cortical flows in polarized *C. elegans* zygotes. We used quantitative analysis of actomyosin network architecture, density, and assembly/disassembly kinetics to calibrate simulations of cortical flows observed in unperturbed wild type embryos. The resulting model predicts changes in the rates and spatial patterns of cortical flow produced by depleting embryos of profilin/PFN-1, formin/CYK-1, and plastin/PLST-1. The success of the model suggests that the variations in cortical flows observed in our experiments are controlled by the variations in architecture of the actin-myosin-plastin network and not by other features (e.g., hydrodynamics). Furthermore, the model provides insights into molecular densities and microscopic rates that are challenging to measure directly *in vivo*.

Cortical flows are driven by a spatial imbalance of contractility, such that some regions contract while others expand. Contraction or expansion is determined by the balance of active forces and internal resistance to deformations within each region. The degree of network connectivity tunes whether each component of the cortex contributes to active forces, internal resistance, or both^1,2,20^. As we decreased the crosslinker density, nucleation rate, and elongation rate in simulations, we observed that the maximum flow speed varied nonmonotonically with the strength of perturbation. Depletions of plastin/PLST-1, formin/CYK-1, and profilin/PFN-1 *in vivo* largely supported these model predictions. Because all of these perturbations decrease the network connectivity, our interpretation of the experiments is that moderate reductions in connectivity lower the resistance to deformation, while strong perturbations produce a catastrophic loss of connectivity (tearing) such that the network can no longer produce and/or transmit forces.

Actin networks tend to resist dilation much more strongly than compression (reviewed in Ref 22) thus it is likely that moderate reductions in connectivity primarily reduce the effective resistance of the posterior cortex to dilation, while stronger losses of connectivity primarily affect the production and transmission of contractile forces across the medial cortex. Because the medial cortex is more highly enriched in F-actin and myosin II than the posterior cortex, it is presumably more highly connected and may therefore be less susceptible to perturbations that reduce connectivity. Thus the nonmonotonic response of cortical flow speeds to reduced network connectivity may be due to the different susceptibilities of posterior and medial cortex to changes in connectivity.

Previous studies used a similar agent-based approach to study the kinetics of contractile ring constriction in *C. elegans* zygotes^31,32^. Like ours, these studies highlight the importance of network composition in driving the kinetics of network contraction, and they identify a non-monotonic dependence of ring constriction speeds on the density of (motor and non-motor) crosslinkers. In the contractile ring, the active motor forces driving constriction are opposed locally by (motor and non-motor) crosslinks; in cortical flows, as discussed above, medial contraction is opposed by posterior resistance to expansion. Thus, while the physics that govern the nonmonotonic dependencies of ring constriction and cortical flows on network architecture and composition may be similar, the spatial variations in network composition and response add a further dimension to the microscopic regulation of cortical flow. Controlled comparisons across these two instances of contractility occurring in the same cell within minutes of one another will likely provide further insight.

A separate analysis of the simulation data based on machine learning and an analytical model suggests that an inhomogeneous distribution of crosslinkers along filaments is needed to generate unbalanced forces that drive the cortical flow^58^. This could account for our observation that the perturbation of the filament elongation rate has more pronounced effects than the other perturbations because the filament length impacts the distribution of crosslinkers in addition to the connectivity. Interestingly, this analysis indicates that the parameters that reproduce properties of unperturbed embryos correspond to the highest flow speed at which the actin density is essentially homogeneous^58^. Thus the unperturbed behavior may reflect selection for both cortical flow and homogeneous structure.

In the present study, we broke the symmetry of the model by introducing a gradient of myosin by hand. In the polarized zygote during mitosis, during mitosis, the gradient of myosin II activity is shaped by the spatial patterns of CDC-42 activity^36,48^, which in turn are thought to self-organize through feedbacks between CDC-42, myosin II and other polarity proteins^36,59^. Incorporating these signaling dynamics and their interplay with mechanics is a natural next step for the model. Parameterizing the model is likely to become more challenging as it grows in complexity. It would be interesting to leverage physics-informed machine learning methods ^60–62^ not only to obtain the parameters but also to compare the forms of interactions in a particle-based model and/or terms in a continuum model.

## Supplementary Movie Legends

**Movie S1.** Plastin/PLST-1 dynamics in mitosis stage *C. elegans* embryo expressing endogenously tagged plastin (PLST-1::GFP). Time compression 1:30.

**Movie S2.** Simulated filament growth and disassembly reproduces dynamic treadmilling.

**Movie S3.** Quasi-steady cortical flows during mitosis in a one-cell C. elegans embryo expressing endogenously tagged myosin II (NMY-2::mKate2). Time compression 1:30.

**Movie S4.** Simulated cortical flows using reference parameters. Only the simulated actin filaments are shown. Time compression 1:30.

**Movie S5.** Comparison of quasi-steady cortical flows during mitosis in one-cell *C. elegans* embryos after weak, moderate and strong depletion of profilin/PFN-1. All embryos express endogenously tagged myosin II (NMY-2::mKate2). Time compression 1:30.

**Movie S6.** Comparison of quasi-steady cortical flows during mitosis in one-cell *C. elegans* embryos after weak, moderate and strong depletion of formin/CYK-1. All embryos express endogenously tagged myosin II (NMY-2::mKate2). Time compression 1:30.

**Movie S7.** Comparison of quasi-steady cortical flows during mitosis in one-cell *C. elegans* embryos after weak, moderate and strong depletion of plastin/PLST-1. All embryos express endogenously tagged myosin II (NMY-2::mKate2). Time compression 1:30.

## Methods

### C. elegans culture and strains

We cultured the *C. elegans* strains listed in Table 1 under standard conditions^63^ at 20-22 °C on 60 mm plates containing OP50 on normal growth medium. Table 1 provides a list of strains used in this study. Unless otherwise specified, strains were provided by the Caenorhabditis Genetics Center, which is funded by the National Center for Research Resources.

**Table 1:** Worm strains used in experiments.

| Strain Name | Description | Genotype | Source |
| --- | --- | --- | --- |
| EM319 | NMY-2::mKate;<br>GFP::UtrophinABD | NMY-2(cp52[NMY-2::mkate2 + LoxP unc-119(+) LoxP]) I; unc-119(ed3) III;mgSi3[cb-UNC-119 (+) GFP::UtrophinABD]II | Ref 8 |
| EM394 | NMY-2::mKate,<br>PLST-1::GFP | plst-1(msn190[plst-1::GFP]) IV; NMY-2(cp52[NMY-2::mkate2 + LoxP unc-119(+) LoxP]) I; | This study |
| EM460 | NMY-2::mKate, CYK-1::GFP | him-8(e1489) IV; NMY-2(cp52[NMY-2::mkate2 + LoxP unc-119(+)<br>LoxP]) I; cyk-1 (knu83 C-terminal GFP, unc-119 (+)); | This study |
| RZB 213 | PLST-1::GFP | plst-1(msn190[plst-1::GFP]) IV; zbIs2[pie-1::Lifeact::mCherry, unc-119(+)]; | Ref 40 |
| EM342 | LifeAct::mCherry, CYK-1::GFP | cyk-1 (knu83 C-terminal GFP, unc-119 (+)); zbIs2[pie-1::Lifeact::mCherry, unc-119(+)]; | This study |

### RNA interference (RNAi)

We used a previously established feeding method^64^ to perform RNAi experiments. We obtained bacteria targeting formin/CYK-1 and profilin/PFN-1 from the Kamath RNAi library^65^. We used bacteria targeting GFP (a gift from Jeremy Nance) to deplete plastin/PLST-1 in a strain expressing endogenously-tagged PLST-1 (PLST-1::GFP). We grew bacteria to log phase in LB medium with 50 ug/ml ampicillin, seeded ∼300 µl of cultured bacteria onto NGM plates with 1 mM IPTG, incubated seeded plates for ∼36 h at room temperature and then stored them at 4 °C for up to ten days before use. Prior to imaging, we transferred L4 larvae to RNAi plates and cultured them at room temperature for different periods of time before imaging (see Table 2 for the ranges of incubation times for each RNAi experiment).

**Table 2:** Details of RNAi experiments.

| Target gene | RNAi strain | Incubation time |
| --- | --- | --- |
| Plastin | GFP-RNAi | 28-40 h |
| Formin | CYK-1 RNAi | 18-30 h |
| Profilin | PFN-1 RNAi | 12-24 h |

We quantified the strength of plastin/PLST-1 depletion by quantifying the density of PLST-1::GFP on the posterior cortex during mitosis in one-cell embryos. We quantified the strength of formin/CYK-1 depletion by quantifying the density of CYK-1::GFP speckles at the two-cell stage on the cortex of AB cells in mitosis. We quantified the strength of profilin/PFN-1 depletion by measuring the speeds of tracked CYK-1::GFP speckles at the two-cell stage on the cortex of AB cells in mitosis. See below for further details.

### Fixation and phalloidin staining

We fixed and stained embryos with phalloidin as previously described^66^. Briefly, we raised embryos to the young adult stage, transferred them into a solution containing 1.5% sodium hypochlorite (Sigma, St. Louis, MO), and 250 mM KOH for 5 minutes to dissolve the worms and release their eggs. We then collected the eggs by centrifugation, rinsed them three times in egg salts, and then fixed them for 15 min at room temperature (RT) in 3% formaldehyde and 0.2% glutaraldehyde (Electron Microscopy Sciences, Hatsfield, PA) in a cytoskeletal stabilization buffer containing 0.1 mg/ml lysolecithin (Sigma), 60 mM Pipes, 25 mM HEPES, 10 mM EGTA, 2 mM MgCl2, and 100 mM dextrose. After fixation, we rinsed embryos three times in PBS with 0.1% Triton-X (PBT) and then stained them for 1 h at RT with Alexa 488 phalloidin (Molecular Probes) at 1 U/200 µl in PBT. Finally, we rinsed embryos in PBS and mounted them on poly-lysine coated coverslips in 90%Glycerol in PBS before imaging.

### Near TIRF microscopy

We mounted embryos for live imaging as previously described^2^ in standard egg salts on 2% agarose pads. For two-color timelapse imaging of myosin II (NMY-2::mKate2) and formin (CYK-1::GFP) or plastin (PLST-1::GFP), we used a Nikon Ti-E inverted microscope equipped with solid state 50-mW 481 and 561 Sapphire lasers (Coherent), a TIRF illuminator, and a Ti-ND6-PFS Perfect Focus unit. A laser merge module (Spectral Applied Research; LMM5) equipped with an acousto-optical tunable filter (AOTF) allowed rapid (1-2 ms) switching between excitation wavelengths. We collected near TIRF images using a CFI Apo 1.45 NA oil immersion TIRF objective onto a PRIME 95B CMOS camera, yielding a pixel size of ∼107 nm. Image acquisition was controlled by Metamorph software.

For fast near TIRF imaging of PLST-1::GFP, we used an Olympus IX50 inverted microscope equipped with an Olympus OMAC two-color TIRF illumination system, a CRISP autofocus module (Applied Scientific Instrumentation), and a 1.45 NA oil immersion TIRF objective. Laser illumination at 488 nm from a 50-mW solid-state Sapphire laser (Coherent) was delivered by fiber optics to the TIRF illuminator. Images were magnified by 1.6x and collected on an Andor iXon3 897 EMCCD camera, yielding a pixel size of 100 nm. Image acquisition was controlled by Andor IQ software.

For all experiments, we chose a laser illumination angle to maximize the signal-to-noise ratio while maintaining approximately even illumination across the field of view and used the same angle for all observations. Further details of the imaging conditions used for specific quantitative analyses are provided below.

### Plastin streak analysis

We collected a sequence of images from embryos expressing PLST-1::GFP on the Olympus TIRF microscope with 30% laser power and 100 ms exposures. We manually identified individual events in which bright streaks of PLST-1::GFP fluorescence could be observed to grow and then disappear. In FIJI, we used maximum intensity projections to identify the path of the growing streak; we used the Straighten plugin (https://imagej.net/plugins/straighten) to extract a linear pixel array of width 5 pixels along the path for each frame of the image sequence and converted the resulting image stack into a kymograph for analysis.

### Bundle size analysis

To analyze bundle sizes, we fixed and stained embryos with phalloidin as described above. We acquired images on a Zeiss LSM 980 microscope equipped with the Airyscan 2 detector. Images were acquired with a 40X 1.4NA oil immersion lens using the MPLX SR-4X mode. For each embryo, we collected Z-stacks of N images with Z steps of 0.16 μm, processed individual frames with Zen Blue 3.0 software using the Airyscan processing feature with default settings, and then projected the stacks to obtain single images for analysis.

We estimated actin bundle sizes from these images as follows. We identified candidate bundles by measuring intensity profiles along linear transects across the posterior cortex and looking for peaks in intensity. For each candidate peak, we measured intensity profiles for transects with different angles passing through the peak position. For each intensity profile with a single peak that was flanked by minima at least 70% below the peak, we computed the integrated intensity between the minima. We used the smallest integrated intensity for each bundle (corresponding to the transect closest to perpendicular) to represent its size. We manually identified the dimmest filament-like feature in the image and took its integrated intensity to represent the value for a single filament; we computed the size of a bundle as the ratio of its integrated intensity to the integrated intensity for a single filament. We used the same procedure for both experimental and simulated images.

### Measuring cortical flow in mitosis

To measure cortical flows in embryos expressing NMY-2::mKate2, we collected data in timelapse mode at 1 frame per second using 30% laser power and 200 msec exposures. We measured axial velocity profiles from these data using quantitative speckle Fluorescence Microscopy (QFSM) software written in MATLAB by the Danuser group (https://github.com/DanuserLab/QFSM)^53,67^. Briefly, we selected a small region outside of the cortex to determine the background level and only considered videos in which the ratio of the signal-to-noise was greater than 2.4 to ensure that the measurements were robust. We then manually set the intensity threshold for the mask that QFSM uses and ran the speckle detection and tracking using standard QFSM settings^67^.

To compute axial flow profiles in wild type and perturbed embryos, we selected 60 frames during mid-late mitosis when flow patterns and speeds show minimal variation. We computed the time-averaged mean speckle velocity within 1 μm bins along the anterior-posterior axis. We used the inflection point in the axial flow profile to define the medial boundary for each embryo. We used the medial boundary position to align flow profiles across multiple embryos.

### Quantifying depletion of plastin, formin, and profilin

For plastin/PLST-1 depletion experiments, we used RNAi to deplete endogenously tagged PLST-1::GFP, and used the background-subtracted mean intensity of PLST-1::GFP measured on the posterior cortex in early mitosis. For formin/CYK-depletion experiments, we used embryos co-expressing NMY-2::mKate2 and CYK-1::GFP. For each embryo, we first imaged NMY-2::mKate2 through mitosis to collect data for measurements of cortical flow as described above. Then in the same embryo, after first division, we collected a sequence of images of CYK-1::GFP at 1 s intervals using 50 ms exposures and 100% laser power, conditions that allow robust detection of single CYK-1 dimers, using the Kilfoil implementation of the Crocker-Grier algorithm for particle detection as previously described^8^. We then used the average density of CYK-1 speckles detected on the cortex of the anterior AB cell over 50 s in mitosis as a measure of CYK-1 depletion.

Finally, for profilin/PFN-1 depletion, we measured cortical flows in one-cell embryos as described above. Then in the same embryos, at the two-cell stage, we imaged CYK-1::GFP in streaming mode with 100% laser power and 50 ms exposures and performed particle tracking as previously described^8^. We selected the subset of trajectories for which log(mean square displacement) vs. log(time) increased with a slope >1.5, consistent with ballistic motion. We confirmed by direct inspection that these trajectories corresponded to rapid directional motion of formin dimers. Because this motion reflects the elongation of single actin filaments^8^, we took the mean speed of these trajectories as an estimate of mean elongation rate, and thus an estimate of the effective strength of profilin depletion. In embryos strongly depleted of profilin, rapid local cortical movements associated with local network tearing, we manually selected a subset of CYK-1 trajectories located within the most stable cortical regions and used these to compute an estimate of the average elongation rate.

### Simulation details

All agent-based simulations were performed with the software Cytosim (https://gitlab.com/f-nedelec/cytosim)^27,43,44^. Filaments were represented as flexible fibers, while crosslinkers and motors were represented as point objects that moved by Langevin dynamics in two dimensions^44^. To model filament treadmilling, we created a customized fiber class that allowed separately tuning assembly and disassembly rates. Each object had two independent “timer” variables: one sets the time at which elongation from the plus end stops, and the other sets the time at which disassembly from the minus end begins.

Above, we present results for a simulation region that is 16 μm x 20 μm; in practice, to minimize discontinuities in molecular densities, we employed a 32 μm x 20 μm region with periodic boundaries in both dimensions and computed statistics from one half (Figure 10). We successively tuned the parameters for individual filament assembly and disassembly, crosslinking, and filament nucleation (and thus density) as described in the Results. To create a myosin gradient, we initialized the system with a higher density of myosin motors in the center (Figure S1) and, throughout the simulation, added myosin motors to the center of the system at a higher rate than to the periphery while removing them without any spatial bias. To reproduce the observed myosin gradient and maximum average flow, we optimized three parameters: the base addition rate of myosin in the system, the addition rate in the center relative to the base addition rate, and the total number of myosin motors. We first varied the base addition rate with the addition rate in the center fixed at 10 times the base addition rate to tune the maximum velocity (Figure S2A) and the myosin densities at the center and edges of the simulation region (Figure S2B). Based on these results, we fixed the base addition rate at 30 s^−1^ and varied the ratio of the center addition rate to the base addition rate (Figure S3); we set this ratio to 6. We then tuned the initial number of myosin motors to reproduce the experimentally observed flow speed.

We summarize the parameters for modeling unperturbed embryos in Table 3 and the perturbations to simulate RNAi experiments in Table 4.

**Table 3:** Simulation parameters and their sources.

|  | Value | Source/citation |
| --- | --- | --- |
| System |  |  |
| Size | 32 $\mu\text{m}$ x 20 $\mu\text{m}$ | Measured from size of posterior in *C. elegans* embryos |
| Boundary Conditions | Periodic | Chosen based on properties of experimental region of interest |
| kT | 0.0042 pN/$\mu\text{m}$ | Thermal energy at 25° C |
| Viscosity | 1 pN s/$\mu\text{m}^2$ | Ref 62 |
| **Actin filaments** |  |  |
| Actin filament rigidity | 0.075 pN/$\mu\text{m}^2$ | Ref 62 |
| Actin filament segmentation | 0.2 $\mu\text{m}$ | Computational parameter, recommended by Cytosim guidelines, Ref 62 |
| Actin filament nucleation rate | Varied, as described in Table 4 | Tuned to match overall network density in simulations |
| Actin filament assembly/disassembly speed | Varied, as described in Table 4 | Ref 8 |
| Assembly timer | 7 + t s, where t is drawn from an exponential distribution with mean 1, corresponding to an average assembly time of 8 s. | Ref 8 |
| Disassembly timer | 7 + t s, where t is drawn from an exponential distribution with mean 1, corresponding to an average disassembly time of 8 s. | Chosen to match measured lifetime of polymerized filaments, Refs 8,45,46 |
| **Crosslinkers** |  |  |
| Binding rate | 60 s<sup>-1</sup> | Optimized to reproduce filament-guided filament assembly |
| Unbinding rate | 0.05 s<sup>-1</sup> | Ref 40 |
| Binding range | 10 nm | Ref 40, 24 |
| Diffusion | 10 $\mu\text{m}^2$ /s | Ref 40 |
| Stiffness | 250 pN/$\mu\text{m}$ | Ref 40, 24 |
| Addition rate | 50 s<sup>-1</sup> |  |
| Removal rate | 50 s<sup>-1</sup> | Set to the addition rate |
| Initial number | Varied, as described in Table 4 | Optimized to reproduce bundle sizes |
| **Motors** |  |  |
| Binding rate | 5 s<sup>-1</sup> | Ref 40 |
| Unbinding rate | 0.15 s<sup>-1</sup> | Ref 40 |
| Binding range | 10 nm | Ref 40, 24 |
| Diffusion | 10 $\mu\text{m}^2$ /s | Ref 40 |
| Stiffness | 250 pN/$\mu\text{m}$ | Ref 40, 24 |
| Unloaded speed | 0.3 $\mu\text{m/s}$ | Measured speed of myosin minifilaments undergoing ballistic motion |
| Stall force | 45 pN | Calculated from Ref 44,52 |
| Base addition rate | 30 $\text{s}^{-1}$ | Optimized to reproduce the maximum flow speed |
| Center addition rate | 180 $\text{s}^{-1}$ | Optimized to reproduce the densities at the center and edges of the simulation |
| Removal rate | 210 $\text{s}^{-1}$ | |
| Initial number | 1,600 | Optimized to reproduce the maximum flow speed |

**Table 4:** Ranges of perturbations for Cytosim simulations.

| Perturbation | Minimum Value | Maximum Value | Unperturbed Value |
| --- | --- | --- | --- |
| Crosslinker number | 445 | 53,400 | 44,500 |
| Actin filament nucleation rate | 12.1 $\text{s}^{-1}$ | 181.5 $\text{s}^{-1}$ | 121 $\text{s}^{-1}$ |
| Actin filament elongation rate | 0.15 $\mu\text{m/s}$ | 2.0 $\mu\text{m/s}$ | 1.5 $\mu\text{m/s}$ |

### Simulated images (pseudoimages)

To create pseudoimages, we estimated the point spread function associated with different labels and imaging conditions (e.g., phalloidin-stained actin filaments visualized with AiryScan microscopy vs. myosin visualized with TIRF). We convolved the locations of components of interest with their corresponding point spread functions. This allowed us to analyze simulations and experiments by identical means.

### Quantifying coherence of motion

To quantify the coherence of motion in the simulations, we computed the correlation length of the velocity component perpendicular to the flow direction (*v_y_*). The data for each condition were from 10 independent simulations of 40 s; we saved frames every 1 s to obtain 40 frames per simulation or 400 frames per condition. In a given frame, we computed *v_y_* for each bound crosslinker and then the product *v_y_*(0)*v_y_*(*d*) for all pairs of bound crosslinkers, where *d* was the distance between the crosslinkers; we binned the distances and averaged *v_y_*(0)*v_y_*(*d*) over 4 blocks of 10 successive frames from each simulation. We fit the correlation function for each block with the form *C*_0_exp(−*d*⁄*d*_0_) to obtain the correlation length (*d*_0_) and the value at *d* = 0(*C*_0_). We report the averages of *d*_0_ and *C*_0_ and their standard deviations (error bars) over the 40 fits (Figures 6, S4, and S5).

## Acknowledgments

This work was supported by National Institutes of Health award number GM 143576 and National Science Foundation award numbers MCB-2201235 and PHY-2317138. The simulations were performed on resources provided by the University of Chicago Research Computing Center.

---

**Figure S1:**
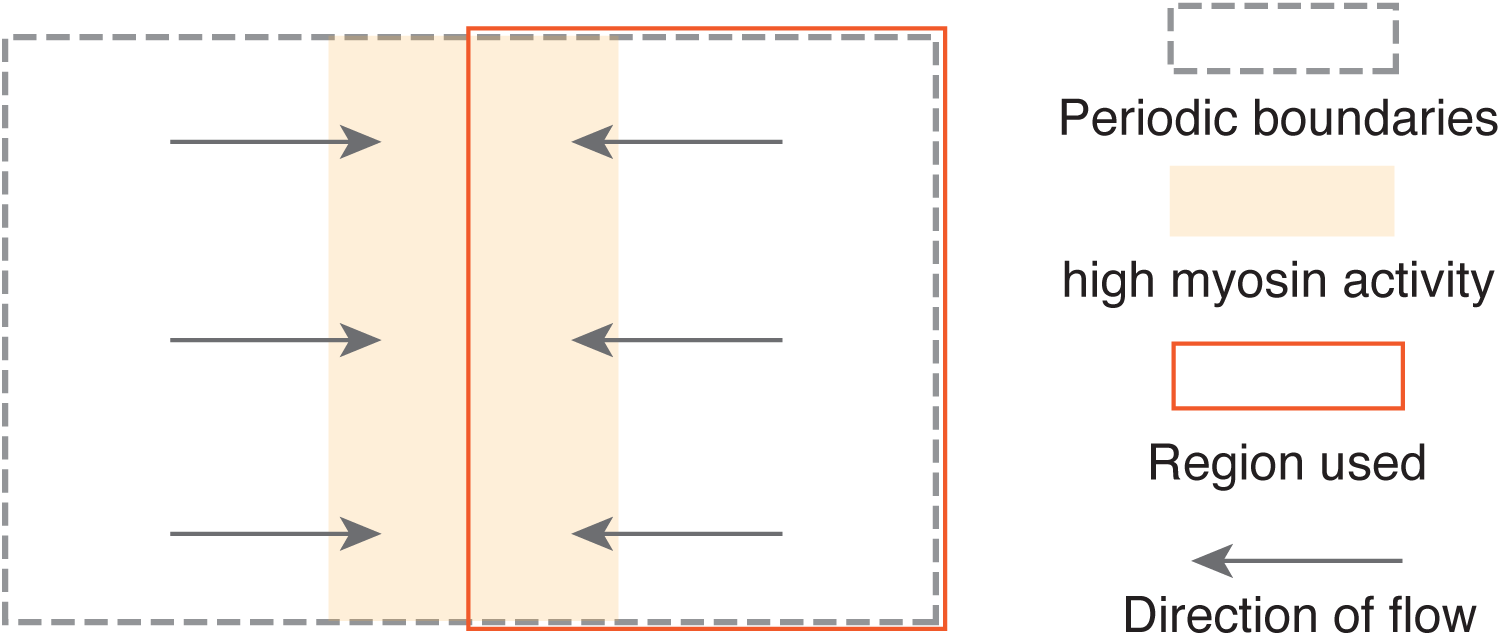
Schematic overview of Cytosim simulations. We performed all simulations on a 32 x 20 μm rectangular domain with periodic boundaries on the x and y axes. Myosin was enriched in the center 10 μm of the simulation, which produced flow towards the center of the space. By symmetry, simulated patterns of flow within the left and right halves of the domain are mirror symmetric about X = 0 and flow speeds vanish at the midline (X = 0) and at the left and right boundaries. We compared simulated flows within the right half domain between X = 0 and X = 16 to our experimental observations.

**Figure S2:**
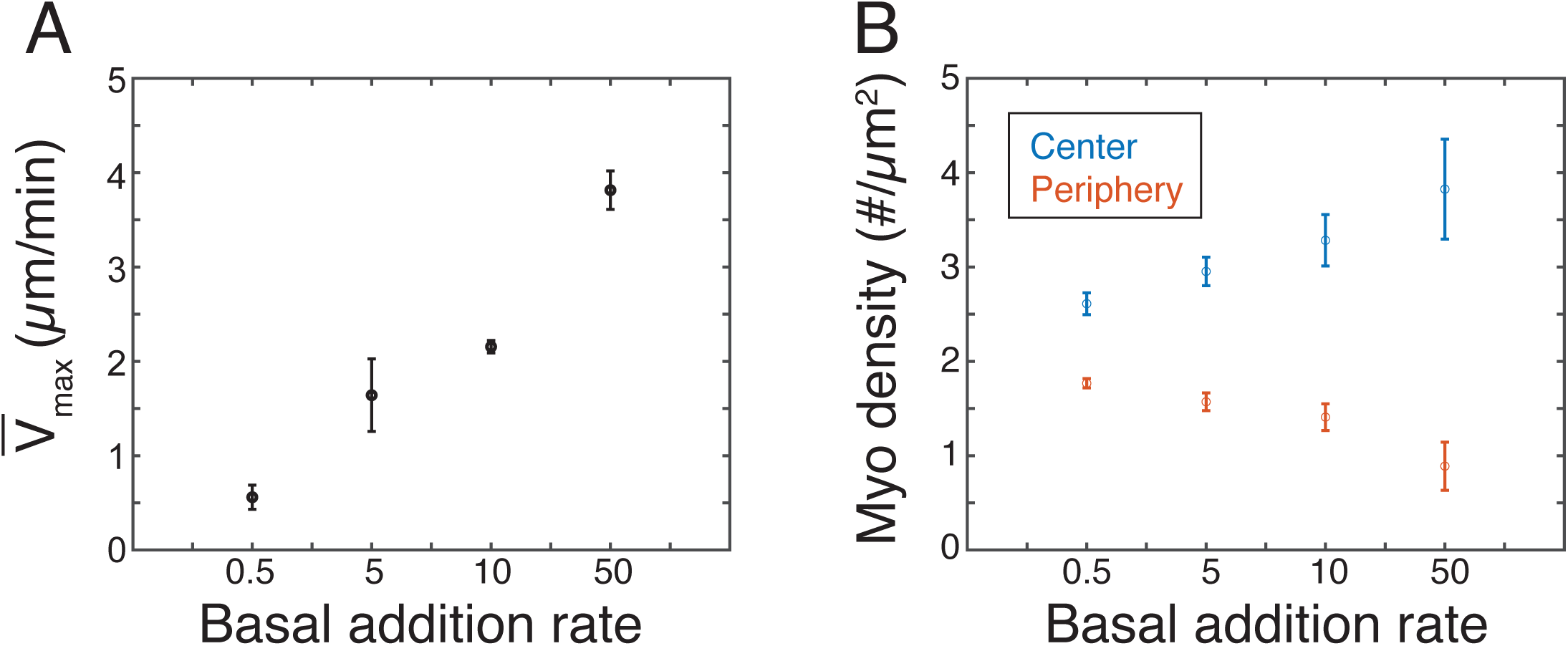
Calibration of the base addition rate of myosin motors. (A) Maximum flow speed increases as base addition rate increases. (B) The difference between the densities of myosin motors at the center (blue) and periphery (red) increases at higher base addition rates. Note that the base addition rate scale is not linear.

**Figure S3:**
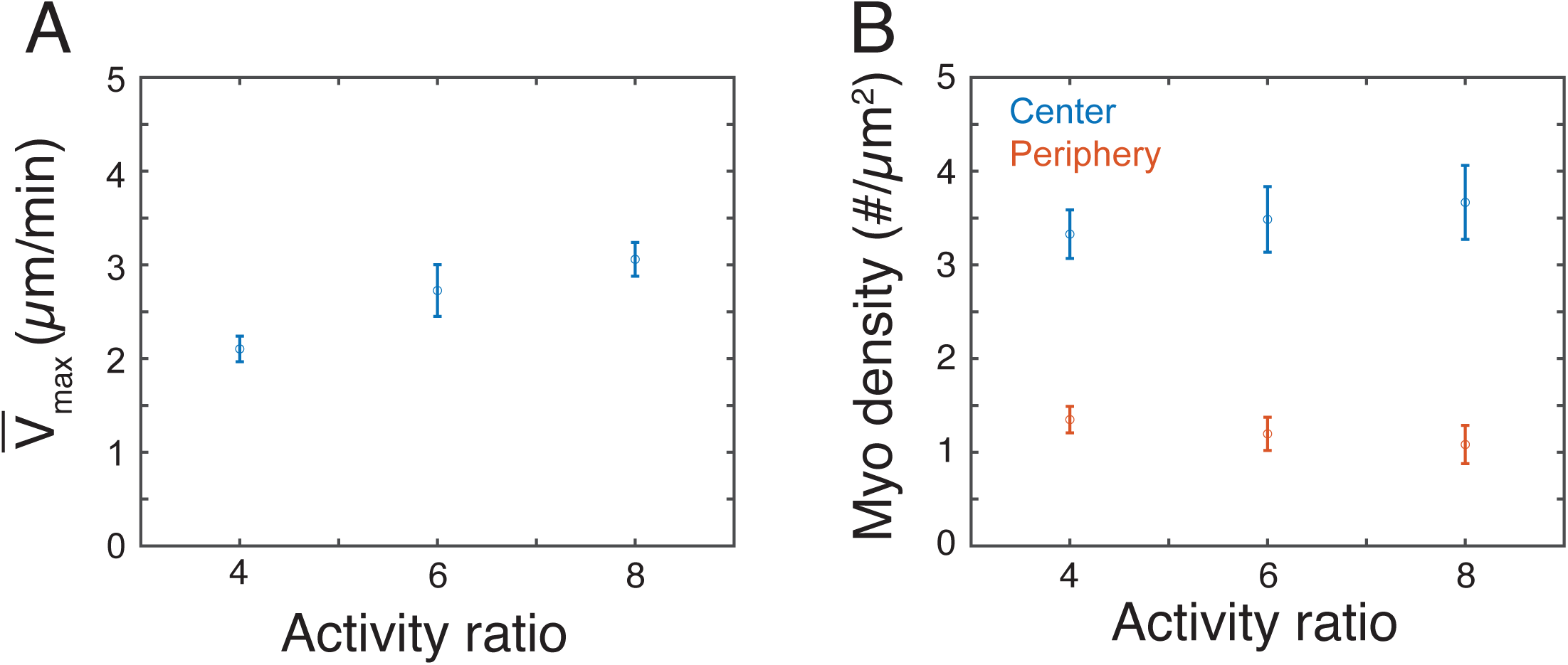
Calibration of the ratio of the center addition rate of myosin to its base addition rate. (A) Maximum flow speed increases with this ratio. (B) The difference between the densities of myosin at the center (blue) and periphery (red) increases with this ratio.

**Figure S4:**
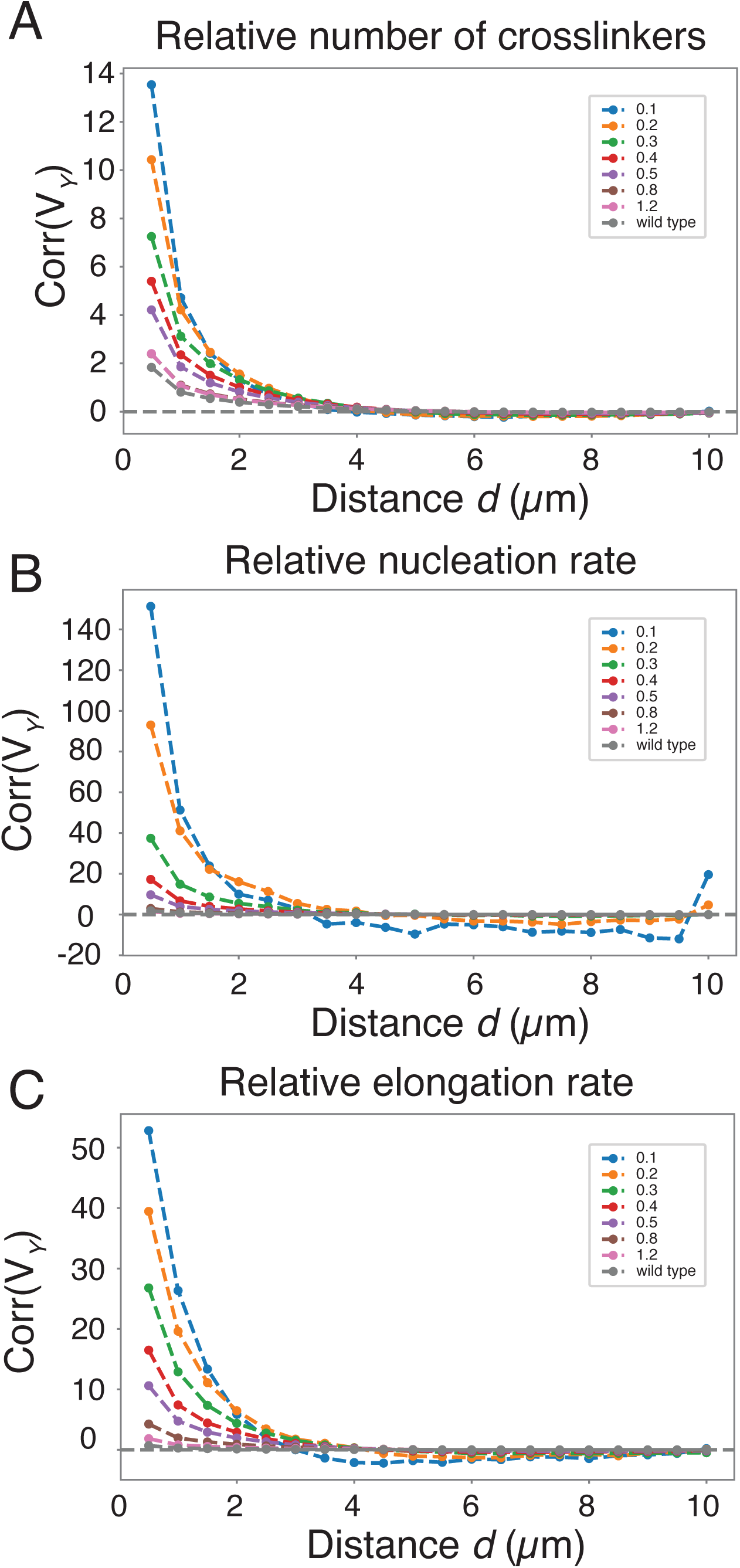
Estimation of correlation length from simulated flows under three different perturbations. The correlation length d_0_ of the correlation function in Figure 6 was determined by fitting the positive values of the correlation function with the form C_0_ = exp(-d⁄d_0_), where d is the distance between crosslinkers and C_0_ is the value of the correlation function at d = 0. Inset legend indicates the different perturbations. Horizontal axis measures the strength of each perturbation as a fraction of the wild type reference value.

**Figure S5:**
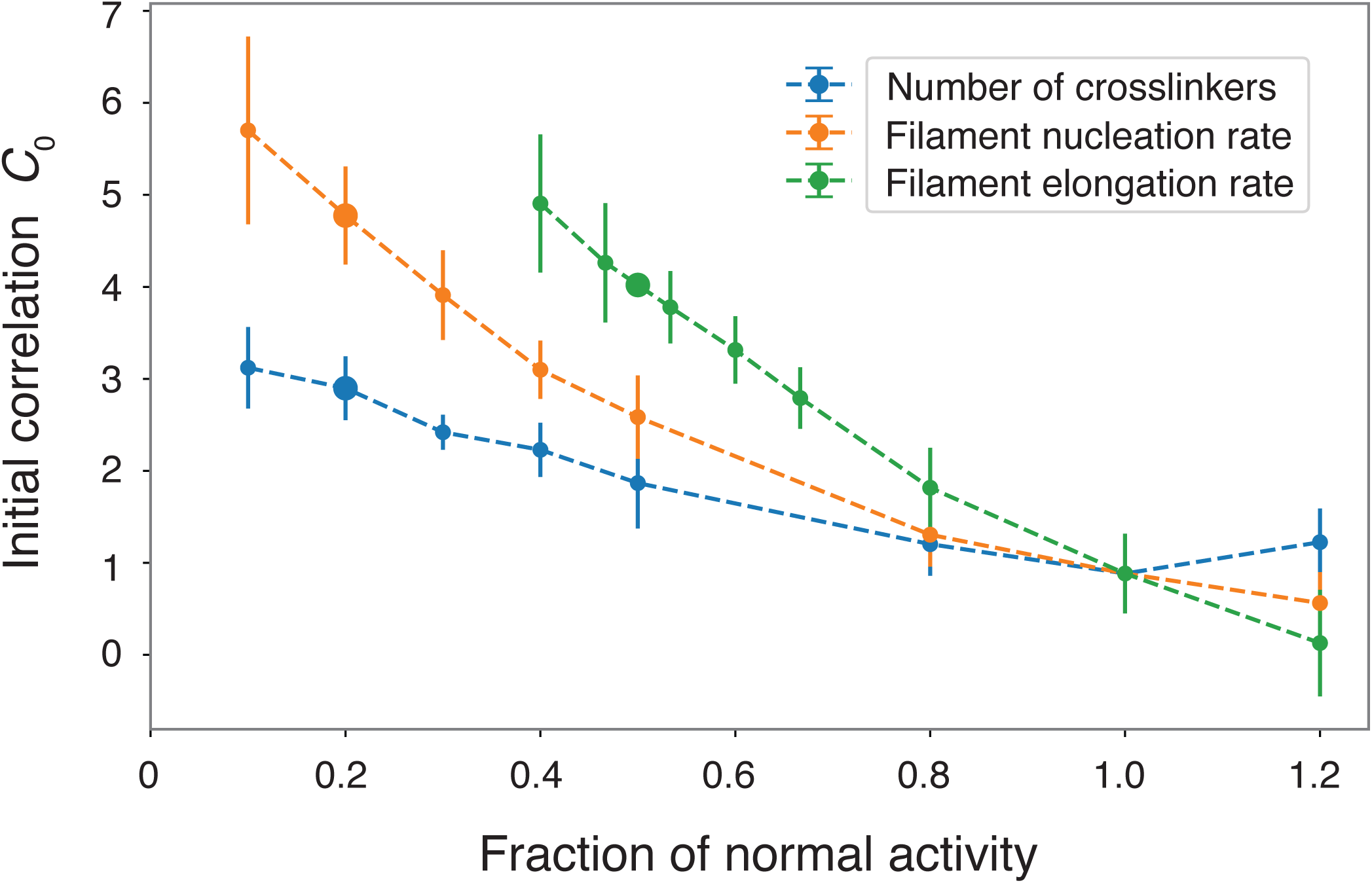
Estimates of initial correlation values C_0_ from fits to the correlation functions shown in Figure S4. Inset legend indicates the different perturbations. Horizontal axis measures the strength of each perturbation as a fraction of the wild type reference value.

## Notes

### Competing Interest Statement

The authors have declared no competing interest.

